# A metagenomics-based survey of the virus diversity in mosquito vectors allows the first detection of Sindbis virus in Burkina Faso

**DOI:** 10.1101/2024.02.02.578537

**Authors:** Didier P. Alexandre Kaboré, Antoni Exbrayat, Floriant Charriat, Dieudonné Diloma Soma, Simon P. Sawadogo, Georges Anicet Ouédraogo, Edouard Tuaillon, Philippe Van de Perre, Thierry Baldet, Roch K. Dabiré, Patricia Gil, Serafin Gutierrez

## Abstract

Mosquito-borne viruses represent a threat to human health worldwide. This taxonomically-diverse group includes numerous viruses that recurrently spread into new regions. Thus, periodic surveillance of the arbovirus diversity in a given region can help optimizing the diagnosis of arboviral infections. Nevertheless, such screenings are rarely carried out, especially in low-income countries. Consequently, case investigation is often limited to a fraction of the arbovirus diversity. This situation probably results in undiagnosed cases. Here, we have explored the diversity of mosquito-borne viruses in two regions of Burkina Faso. To this end, we have screened mosquitoes collected along three years in six urban and rural areas using untargeted metagenomics. The analysis focused on two mosquito species, *Aedes aegypti* and *Culex quinquefasciatus*, considered among the main vectors of arboviruses worldwide. The screening detected *Sindbis virus* (SINV, *Togaviridae*) for the first time in Burkina Faso. This zoonotic arbovirus has spread from Africa into Europe. SINV causes periodic outbreaks in Europe but its distribution and epidemiology in Africa remains largely unstudied. SINV was detected in one of the six areas of the study, and at a single year. Detection was validated with isolation in cell culture. SINV was only detected in *Cx. quinquefasciatus*, thus extending the list of potential vectors of SINV in nature. SINV infection rate in mosquitoes was similar to those observed in European regions that experience SINV outbreaks. A phylogenetic analysis placed the nearly-full genome within a cluster of Central African strains of lineage I. This cluster is supposedly at the origin of the SINV strains introduced into Europe. Thus, West Africa should also be considered as a potential source of the European SINV strains. Our results call for studies on the prevalence of SINV infections in the region to estimate disease burden and the interest of SINV diagnostic in case investigation.

**Author summary:** Mosquito-borne viruses are responsible for millions of cases worldwide every year. Moreover, they have repeatedly shown an ability to spread over large distances. Thus, periodic surveys of the arbovirus diversity in a given region can help to define the diagnostic tests to use during case investigation. However, comprehensive surveys are rarely carried out, especially in low-income countries. Here, the arbovirus diversity was assessed in two main mosquito vectors in Burkina Faso using untargeted metagenomics. This screening identified *Sindbis virus* (SINV), a zoonotic arbovirus, for the first time in Burkina Faso. Moreover, SINV was found in nature for the first time in *Culex quinquefasciatus*, a main mosquito vector of several pathogens and with a cosmopolitan distribution. SINV leads to periodic outbreaks mainly in Europe. Despite a likely African origin, its distribution and epidemiology in Africa remains largely unstudied. The SINV sequence from Burkina Faso felt within the cluster of Central African strains thought to be at the origin of the European SINV strains. Thus, our results indicate that West Africa should be considered as another potential source of the SINV introductions in Europe. Further studies are required to characterize SINV epidemiology in Burkina Faso and the West African region.

## Introduction

Emergences and re-emergences of mosquito-borne viral diseases have become more frequent and intense over the last decades [1]. This trend is probably due to a combination of factors, including global transport, urbanisation and climate change [1,2]. The increase in incidence and geographical distribution has been observed for different virus species representative of the wide taxonomic diversity of mosquito-borne viruses [3,4]. To date, this diversity potentially includes 245 virus species of RNA viruses, with most species belonging to the families *Flaviviridae* and *Togaviridae*, and the order *Bunyavirales* [5]. Moreover, this diversity includes a variety of life cycles, differing in the species of mosquito vectors and vertebrate hosts [2]. Not surprisingly, different epidemiological situations are associated to these viruses. For example, dengue virus stands out in case number, with hundreds of thousands of cases per year globally [6]. On the other hand, numerous lesser-known viruses often lead to outbreaks limited in case number, geographical distribution or duration [7].

There is currently no vaccine or antiviral drug available against most of mosquito-borne viruses [8–10]. Prevention and control of arboviruses therefore mainly rely on surveillance. In case of virus detection, mosquito management and public awareness campaigns are usually implemented to, among others, reduce exposure to infected vectors [11]. Nevertheless, the surveillance of mosquito-borne viruses often faces several challenges. One of these challenges is the large diversity of arbovirus species harboured in certain regions [7]. For example, several tenths of mosquito-borne viruses can be found in many African countries [7]. This diversity renders complex the implementation of diagnostic tests for all the viruses present in the region due to, among others, test availability and cost [12]. Another challenge is the potential changes in the diversity of mosquito-borne viruses in a given region along time. Many of these viruses have periodically spread over large distances and colonize new territories. Such long-distance spread is mainly driven by infected vertebrate hosts, like humans or migratory birds [13–17]. If virus spread into non-endemic regions leads to outbreaks, the latter can go unidentified because infections often lead to non-specific signs and symptoms.

Thus, the diversity of mosquito-borne viruses can be high and change over time in certain regions. However, routine diagnostic of potential infections is usually limited to a few viruses with a major impact on public health, like dengue virus, even in countries in which arbovirus diversity is high [18]. This situation can lead to diagnostic failure. While specific treatments for arboviral infections are currently unavailable, a prompt diagnosis can prevent unnecessary diagnostic testing and treatment, as well as provide diagnostic and prognostic information to patients. Moreover, the resulting lack of data on disease prevalence hinders the design and implementation of adequate surveillance and control strategies. Periodic surveys of arbovirus diversity could limit these problems through updated reports on the viruses actively transmitted in a given region. However, comprehensive explorations of the arbovirus diversity are rare, especially in low-income countries [18].

Here, we aimed to characterize the arbovirus diversity in two regions of Burkina Faso. This West-African country is highly connected to other regions in Africa and other continents through human and wildlife movement [19]. Several mosquito-borne viruses have been previously detected in Burkina Faso [20–22]. Nevertheless, studies have focused on a limited number of these viruses. Here, arbovirus diversity was analyzed in a mosquito collection, a classic approach in the surveillance of mosquito-borne viruses [23]. Two mosquito species were targeted, *Aedes aegypti* and *Culex quinquefasciatus*, both considered among the main vectors of arboviruses worldwide. These mosquitoes differ in the arbovirus diversity they transmit [24]. To further facilitate a comprehensive exploration of viral diversity, environments with different levels of human activity, and thus likely to harbour different viruses, were sampled for three consecutive years. Previously, we had screened this mosquito collection for several mosquito-borne viruses - including dengue virus, Zika virus, West Nile virus and chikungunya virus - using RT-PCR [20,21].

In this study, we have used shotgun metagenomics to screen the mosquito collection. Metagenomics is an untargeted diagnostic approach that allows a comprehensive exploration of the virus diversity in a sample and thus avoids the use of a multitude of targeted diagnostic tests [25]. Our approach allowed the first detection of *Sindbis virus* (SINV; family *Togaviridae*, genus *Alphavirus*) in Burkina Faso. The cycle of this zoonotic arbovirus mainly involves mosquitoes and wild birds [26]. SINV can infect humans through mosquito bites. The main symptoms include fever, rash and arthralgia [26,27]. An important proportion of clinical cases lead to joint symptoms over months or years [28]. Nevertheless, most infections are subclinical, a situation leading to a large rate of undiagnosed infections [26]. To date, periodic outbreaks of SINV infections have only been reported in Northern Europe. The limited geographical extent of those outbreaks contrasts the quasi-global distribution of SINV, with reports from Africa, Asia, Australia and Europe [26]. This epidemiological pattern can be partly explained by virus genetics. SINV strains are classified into six genotypes with a limited or no overlap in geographical range [26]. Genotype 1 is the only genotype detected in Europe and Africa, the regions in which outbreaks have been reported. This situation thus suggests a link between SINV genotype and epidemiological risk. Moreover, European strains are the result of a few introductions from Africa, probably through bird migration [16]. SINV strains from Central Africa have been incriminated as the potential ancestors of the European strains [16]. However, the geographical source of European strains remains to be further explored because SINV sequences are only available from a limited number of African regions. Moreover, recent reports suggest that SINV is either spreading into new African regions or endemic in a larger area than previously thought [29,30]. Thus, the paucity of data on SINV hampers to fully evaluate its distribution in Africa and routes of spread.

## Methods

### Collection sites and mosquito sampling

Collection sites have been previously described [31]. Mosquitoes were collected in two regions of Burkina Faso, the Hauts-Bassins and South-West regions (Figure 1). These regions have an average annual rainfall of 1 200 mm. The climate is tropical with two seasons: a rainy season from June to September and a dry season from October to May. The vegetation is mainly composed of tree or wooded savannahs in the Hauts-Bassins region and the South-West regions respectively. Urban and rural zones were sampled in each region. In the Hauts-Bassins region, three zones were sampled, including one urban zone (Urban 1 zone) and two rural zones (Rural 1 and Rural 2 zones). The Urban 1 zone comprised three collection sites in the town of Bobo-Dioulasso, the second largest city in Burkina Faso (904 920 habitants and 13 680 Ha). The Rural 1 zone comprised four sites in a rural area dominated by rice fields 30 km to the north of Bobo-Dioulasso. The collection sites of the Rural 2 zone were situated in two forested areas situated 18 km far away from Bobo-Dioulasso (Nasso and Dinderesso forests). In the South-West region, three zones were distributed over a transect linking two towns, Diébougou (25 688 habitants) and Gaoua (45 284 habitants). Urban 2 and 3 zones comprised a site in either Diébougou or Gaoua, respectively. Moreover, the third zone, Rural 3 zone, included four sites situated in rural areas along the road linking those two cities.

**Figure 1.**
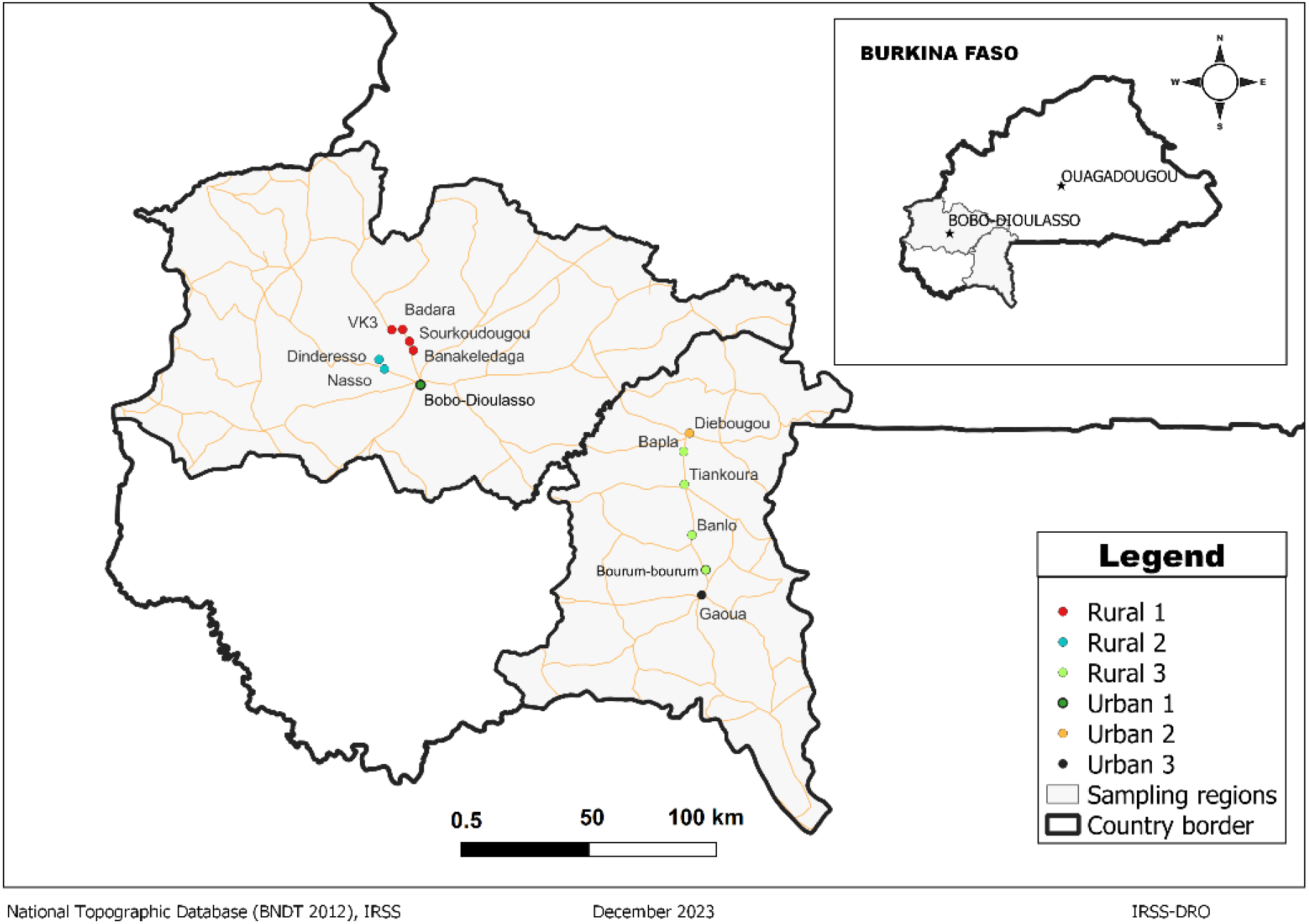
Sites of mosquito collection. Dots indicate collection sites. Dot colour stands for sampling zone (Rural and Urban zones). The inner panel shows the positions of the capital city (Ouagadougou) and the two sampling regions in the country.

Mosquito sampling has been previously described in detail [31]. Briefly, mosquito collection was carried out in all sites in three periods: August/September 2019, June/October 2020 and May/June 2021. Collection of adult mosquitoes was carried out over two successive days in each locality. Three trapping methods were used: the double net tent, the BG-Sentinel traps and the prokopack aspiration. Individual mosquitoes were identified morphologically using identification keys [32,33] on an ice-cold platform to avoid viral genome degradation, then stored at −80°C for subsequent analyses.

### Mosquito pooling and isolation of nucleic acids

Non-blood-engorged females were pooled by date and collection zone (Table 1). A total of 60 pools of *Cx. quinquefasciatus* (1629 females) and 47 pools of *Ae. aegypti* (1356 females) were generated (Table 1). Details per species, area and year on the number of individuals and pools can be found in Table 1. In 2019, mosquito catches were low and pools could only be generated for the Rural 1 and Urban 1 zones.

**Table 1.** Number of females of *Aedes aegypti* or *Culex quinquefasciatus* per collection zone and year, and the resulting pools and libraries for Illumina sequencing. Totals per collection zone appear in bold. The last row provides the totals for all zones together.

| zone/year | *Aedes aegypti* |  |  | *Culex quinquefasciatus* |  |  |
| --- | --- | --- | --- | --- | --- | --- |
|  | Females | Pools | Libraries | Females | Pools | Libraries |
| **Rural 1** | **166** | **6** | **6** | **325** | **12** | **7** |
| 2019 | 90 | 3 | 3 | 17 | 1 | 1 |
| 2020 | 51 | 2 | 2 | 172 | 6 | 3 |
| 2021 | 25 | 1 | 1 | 136 | 5 | 3 |
| **Rural 2** | **47** | **2** | **2** | **158** | **6** | **4** |
| 2019 | 0 | 0 | 0 | 0 | 0 | 0 |
| 2020 | 35 | 1 | 1 | 81 | 3 | 2 |
| 2021 | 12 | 1 | 1 | 77 | 3 | 2 |
| **Rural 3** | **79** | **3** | **3** | **101** | **4** | **4** |
| 2019 | 0 | 0 | 0 | 0 | 0 | 0 |
| 2020 | 52 | 2 | 2 | 44 | 2 | 2 |
| 2021 | 27 | 1 | 1 | 57 | 2 | 2 |
| **Urban 1** | **609** | **21** | **11** | **616** | **22** | **10** |
| 2019 | 57 | 2 | 2 | 77 | 3 | 2 |
| 2020 | 420 | 14 | 7 | 397 | 14 | 5 |
| 2021 | 132 | 5 | 2 | 142 | 5 | 3 |
| **Urban 2** | **278** | **9** | **6** | **195** | **7** | **4** |
| 2019 | 0 | 0 | 0 | 0 | 0 | 0 |
| 2020 | 184 | 6 | 3 | 83 | 3 | 2 |
| 2021 | 94 | 3 | 3 | 112 | 4 | 2 |
| **Urban 3** | **177** | **6** | **4** | **234** | **9** | **5** |
| 2019 | 0 | 0 | 0 | 0 | 0 | 0 |
| 2020 | 121 | 4 | 2 | 48 | 2 | 2 |
| 2021 | 56 | 2 | 2 | 186 | 7 | 3 |
| **TOTAL** | **1356** | **47** | **32** | **1629** | **60** | **34** |

Each pool was processed to obtain nucleic-acid isolations enriched in nuclease-protected molecules as previously described [34]. Briefly, mosquitoes were homogenized in 500 μl ice-cold 1X Phosphate Buffered Saline buffer with two ice-cold steel bearing balls (3 mm diameter, LOUDET) using a TissueLyser II (Qiagen) and clarified through centrifugation. A 150-μl aliquot of clarified homogenates was digested with a nuclease cocktail consisting of 20 U/L of Exonuclease I (Thermofisher), 5 U/L of RNase I (Thermofisher), 25 U/L of Benzonase (Merck Chemical) and 20 U/L of turbo DNase (Ambion). Then, nucleic acids in the resulting suspension were isolated using the Nucleospin RNA virus kit (Macherey Nagel) according to the manufacturer’s protocol with modifications. More specifically, 20 μl of proteinase K at 20 mg/ml (Macherey Nagel) were added per reaction in the RAV1 buffer.

Moreover, the RNA carrier was replaced in RAV1 buffer by 5 μg of linear acrylamide (Ambion). Nucleic acids were eluted in 50 μl of RNase-free water and stored at −80°C. The quality and quantity of the RNA was estimated using capillary electrophoresis (2100 Bioanalyzer, Thermofisher). In addition, a second isolation per pool was carried out as described above but without the nuclease treatment. The resulting total-RNA sample was used for virus detection with RT-qPCR.

### Library preparation and Illumina sequencing

A custom library preparation for Illumina sequencing was conducted from nucleic-acid suspensions enriched in nuclease-protected molecules as described [35]. Briefly, cDNA was generated in a reverse transcription reaction using the RevertAid First Strand cDNA synthesis kit (ThermoScientific) and the 454-E-8N primers. Double stranded DNA (dsDNA) was then generated using the Klenow fragment polymerase (Fisher Scientific) and the 454-E-8N primers. The dsDNA fragments were amplified in a PCR reaction with the 454-E primer and the Phusion High-Fidelity DNA Polymerase kit (Fisher Scientific). Amplicons were purified with the NucleoSpin gel and PCR Clean-up (Macherey-Nagel).

Then, adapters were ligated to the amplicons in a PCR reaction using the P5 and P7-bearing adapter primers and the Phusion High-Fidelity DNA Polymerase kit (Thermo Scientific). Amplicons were sized and purified using an AmpurXP magnetic bead capture (Agencourt). The size of the resulting libraries (expected 500-600 bp) was validated in a capillary electrophoresis (Agilent 2100 Bioanalyzer, Agilent Technologies). Library concentration was estimated with the Library Quantification kit (Takara Bio) according to the manufacturer’s protocol. All libraries were pooled together in similar concentrations and the resulting sample was sequenced in the same run with a HiSeq 2500 sequencer (Illumina; 250-bp paired reads) using specific sequencing primers [35].

### Virus detection from Illumina reads

The bioinformatic analysis of reads was carried out with the Snakevir pipeline [35]. Snakevir and the associated documentation is freely available at https://github.com/FlorianCHA/snakevir. Briefly, removal of adapter and low-quality sequences from reads was done with Cutadapt 1.6 [36]. Then, rRNA-derived reads were removed from the dataset. The identification of rRNA-derived reads was done with a mapping with BWA 0.7.15 [37] against rRNA sequences (SILVA bacterial bases: SSURefNr99 and LSURef; 18/01/2017; SILVA dipteran base, release 132) [38]. Then, reads from all libraries were pooled and used as input in a de-novo assembly to generate a non-redundant set of contigs using Megahit v1.1.2 [39]. The contigs and non-assembled reads were used in a second *de-novo* assembly with CAP3 [40]. The resulting metagenome was screened for virus-derived contigs in a homology search using Diamond v2.1.8 [41] (e-value cutoff = 10^−3^) against the NCBI nr database (May 2021). A further search for SINV-like contigs was done through an assembly of the virus-like contigs that mapped on a SINV genome (Accession OK644705) in Geneious 10.2.6.

### PCR detection of SINV and Sanger sequencing

A SINV-specific real-time RT qPCR [29] was used to screen the 60 pools of *Cx. quinquefasciatus* mosquitoes for the presence of SINV nucleic acids. RT-qPCR was performed using the Luna Universal One-Step RT-qPCR Kit (Biolaps) according to the manufacturer protocol on a LightCycler real-time PCR thermocycler (Roche). The analysis of the infection rate based on PCR data was done with the *binGroup* package [42] in R software.

SANGER sequencing was performed on PCR amplicons covering gaps between SINV contigs. Amplicons were obtained using a set of primers defined on contig sequences (Table S1) with the PCR Kit phusion hight fidelity (Thermofisher) following manufacturer’s instructions.

### Phylogenetic analysis

Multiple alignment of sequences encoding both the nonstructural protein (nsP) and the structural protein (sP) of SINV-Burkina Faso and 27 sequences from SINV genotype 1 was performed with MEGA version 11 software [43]. The sequences were aligned using the Clustal W multiple alignment algorithm [44]. A phylogenetic tree was constructed using the maximum-likelihood method and a General Time Reversible model [45] with 1 000 bootstrap replicates using MEGA version 11 Software.

### Virus isolation

Homogenates of the two mosquito pools in which SINV was detected were used as inoculum for virus isolation in Vero African green monkey cells (Vero ATCC CCL81). For each pool, a volume of 100 μl of homogenate was inoculated on in a T-25 cell culture flask at 37°C in a 5% CO2 atmosphere for one hour and half. Then the supernatant was replaced by 5 ml of MEM medium supplemented with 10% FBS, 200mM L-glutamine, 1% penicillin-streptomycin, 50μg/ml Gentamycin. Cells were incubated at 37°C in a 5% CO2 atmosphere and examined for cytopathic effects daily. Supernatant (150 μl) from the first passage (five days after inoculation) was analyzed for the presence of SINV nucleic acids using the SINV RT-qPCR described above [29].

## Results

We used shotgun sequencing of nuclease-protected nucleic acids to investigate the virus diversity in populations of *Cx. quinquefasciatus* and *Ae. aegypti* from two regions of Burkina Faso. A total of 66 libraries were generated from 2985 mosquitoes and sequenced. Differences in the size of mosquito collections led to differences in the number of libraries per region (40 and 26 libraries from the Hauts-Bassins and South-West regions, respectively; Table 1). The number of libraries was similar between mosquito species but with differences in the number of mosquitoes per library (median number of mosquitoes per library = 51.5 and 31.5 for *Cx. quinquefasciatus* and *Ae. Aegypti*, respectively; Table 1).

Sequencing generated a total of 216 million reads and 3.34 million reads per library on average (min/max = 0.8/7.65 million reads per library). Total read number was similar between mosquito species (1.07 and 1.09 million reads for *Cx. quinquefasciatus* and *Ae. aegypti* respectively). After bioinformatic processing of reads, we identified 1412 virus-like contigs (mean/min/max contig length = 1.14/0.22/28.36 Kb). Virus-like contigs were associated to 103 virus species and mostly to arthropod-specific viruses (85% of the virus species). The description of the full virus diversity is out of the scope of this study and was not further explored here.

A mosquito-borne zoonotic virus was detected. This virus was SINV. A total of twelve contigs were found associated to SINV sequences with percent identities at the amino acid level above 90%. The SINV-like contigs were found in a single library. The library derived from two pools of *Cx. quinquefasciatus* collected in Rural 1 zone in 2020 (Figure 1). The SINV contigs covered 84% of the genome (9 712 bp). The gaps between the contigs were determined with Sanger sequencing. A nearly-full genome (11 244 bp) was obtained and submitted to GenBank (Accession OR659083). A phylogenetic analysis clustered the SINV sequence within genotype 1, the genotype prevalent in Africa and Europe and behind most outbreaks of SINV infection (Figure 2). Moreover, the sequence from Burkina Faso felt within a cluster of sequences, most from Central Africa, that has been shown to be at the origin of the SINV genotypes that have colonized Europe [16].

**Figure 2:**
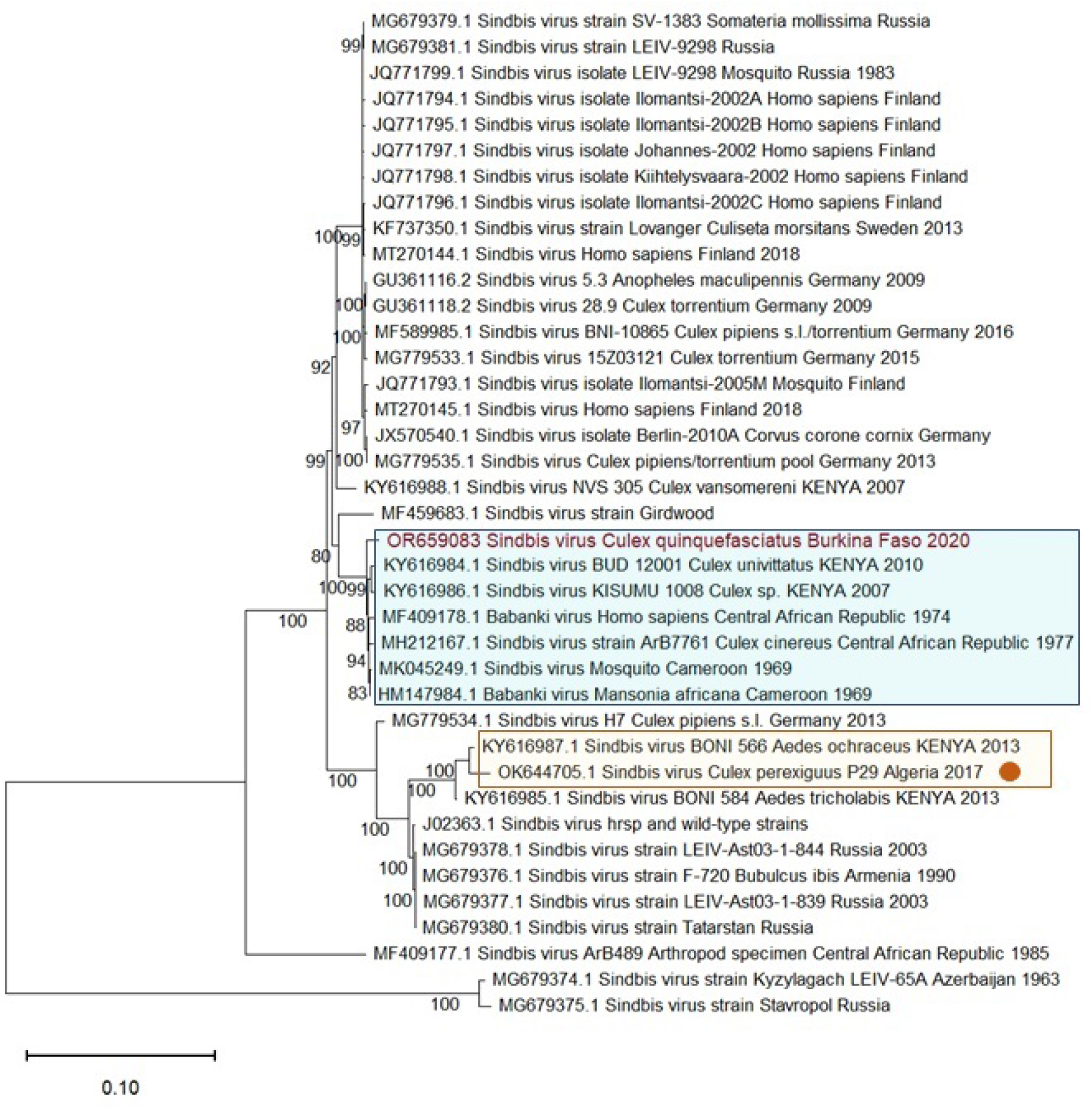
Maximum-likelihood phylogenetic tree of nucleotide sequences from SINV genotype I. Sequences are concatenated open reading frames. The sequence from Burkina Faso appears in red. The sequence from Algeria, from a previous study, appears with an orange circle after the name. Two clusters appear underlined with coloured rectangles: the cluster including the sequence from Burkina Faso (blue rectangle) and that including the sequence from Algeria (orange rectangle). The scale bar represents the number of nucleotide substitutions per site. GenBank accession numbers of the sequences are provided on the branch names.

To estimate the infection rate in mosquito populations, we screened the 60 pools of *Cx. quinquefasciatus* (1629 mosquitoes in total) for SINV nucleic acids with RT-PCR, a technique considered more sensitive than metagenomics. Although *Cx. quinquefasciatus* has not been shown to be infected by SINV in nature, we restricted the screening to this mosquito species because the main SINV vectors usually belong to the *Culex* genus [7,16,29]. Two pools were found positive for SINV nucleic acids. These two pools had been used to generate the SINV-positive library. To validate that the detection of SINV nucleic acids arose from a viable virus, we attempted to isolate the virus in cell culture from the two SINV-positive pools. Cytopathological effects were observed in the cultures inoculated with the pool with the highest virus concentration as shown with RT-qPCR. SINV isolation was confirmed through PCR on culture supernatant. We then inferred the SINV infection rate among *Cx. quinquefasciatus* females in Rural 1 zone in 2020 using the RT-qPCR data. The maximum-likelihood estimate of the infection rate was 1.25% (95% confidence intervals = 0.24%/4.26%).

## Discussion

Our working hypothesis postulated that shotgun metagenomics can be an alternative to targeted diagnostic in surveys of the full diversity of mosquito-borne viruses in a given region. Our metagenomics-based survey allowed to detect SINV, a zoonotic virus, in a recent collection of mosquito vectors. To our knowledge, SINV has not been previously detected in Burkina Faso. SINV detection was validated through virus isolation and PCR from non-blood-engorged females. Previously, a PCR-based survey on the same samples did not detect six mosquito-borne viruses potentially present in the country (dengue virus, Zika virus, chikungunya virus, West Nile virus and Usutu virus) [20,21], just like found here in the metagenomics survey. Comparison of the surveys using either PCR or metagenomics suggests that viral metagenomics on mosquitoes is a suitable tool for virus surveillance in regions with a high viral diversity, as shown by others [46,47]. Nevertheless, metagenomics is yet highly demanding in both financial and technical terms [48]. These constraints should be weighed against the benefits of identifying the circulation of a new pathogen for case diagnostic and patient treatment. Here, we could not determine potential benefits because the prevalence of SINV infections in humans has not yet been studied in Burkina Faso. This is a main limitation of our study. In fact, studies on the prevalence of SINV infections in humans in African countries are scarce [49,50]. The only recent report found a 12.7% incidence in patients with acute febrile disease of unknown cause found in South Africa in 2019 and 2020 [49]. Future studies on the epidemiology of SINV infections in Burkina Faso are needed to assess its potential impact on human health. Those studies can also facilitate to estimate the benefits of the metagenomics-based survey.

This first robust detection of SINV in West Africa could be the result of either a recent introduction into the region, or an endemic situation that had not yet been revealed. This question also applies to the first detection of SINV in Northwest Africa recently [29]. An undetected endemicity in Burkina Faso seems plausible given the limited number of studies on SINV and its large geographical distribution in Africa [15]. However, a recent spread of the virus cannot be ruled out because SINV dispersal has been observed both within and between continents [52,16]. Longitudinal surveys of SINV prevalence and genetics, including retrospective analyses of sample collections, could help to determine whether the same genotype has been present along years, thus supporting endemicity [53].

The phylogenetic analysis provided two interesting results concerning geographical spread of SINV. First, the genomic sequence from Burkina Faso felt within the cluster of sequences from Central and Eastern Africa that are supposedly at the origin of the European strains [16]. Thus, West Africa should also be considered among the potential geographical sources of the European strains. Given the likely importance of bird migration in SINV dispersal, studies on the prevalence and genetics of SINV infections in both sedentary and migratory birds in West Africa could further clarify the potential of the region as a source for SINV spread into Europe. Secondly, the genotype recently identified in Northwest Africa [29] felt in a cluster different than that including the sequence from Burkina Faso (Fig. 2). Hence, if a recent SINV spread has taken place into the two African regions, then two distinct dispersals have taken place because the genotypes incriminated are different. Moreover, our results further support that a diversity of SINV genotypes currently circulate in different African regions, calling for more studies on SINV diversity and geographical range in Africa.

SINV was detected in a rural area, in the vicinity of the second largest city in Burkina Faso, and only in 2020. The detection in a rural area is in accordance with the SINV enzootic cycle involving wild birds and ornithophilic mosquitoes, mainly of the *Culex* genus. In this study, the limited number of *Cx. quinquefasciatus* mosquitoes per year and zone (116 females on average) hampered a robust quantitative analysis of SINV prevalence over years. Despite this limitation, two results support the need for future assessment of the potential epidemiological risk of SINV in Burkina Faso. First, we detected potential SINV infection of *Cx. quinquefasciatus* in nature for the first time. Previously, SINV infection of *Cx. quinquefasciatus* has only been observed in laboratory experiments with a long-established colony. This mosquito is a main vector of several mosquito-borne pathogens, including viruses (*e*.*g*., West Nile virus and St. Louis encephalitis virus) and filarial nematodes in subtropical and tropical areas worldwide. Contrary to *Cx. univitattus*, considered the main SINV vector in most African countries and usually found in rural environments [7,16], *Cx. quinquefasciatus* is abundant in both urban and rural areas in Burkina Faso [31]. Moreover, this mosquito, although mainly ornithophilic, readily feeds on humans [54]. SINV has been isolated from the saliva of experimentally-infected females, strongly suggesting that *Cx. quinquefasciatus* could be a SINV vector [55]. Hence, if its role as SINV vector in nature is validated, *Cx. quinquefasciatus* has several features that should increase the risk of spillover from an enzootic cycle to humans. The second result related to epidemiological risk is the SINV prevalence in *Cx. quinquefasciatus* populations. The infection rate in the area and year of detection was within the same order of magnitude as estimates from populations of *Culex* mosquitoes in Europe, including infection rates from situations with outbreaks [56,57]. The observed infection rate could thus be compatible with a potential spillover to human populations.

Burkina Faso, like many African countries, harbours numerous and taxonomically-diverse mosquito-borne viruses that can have an impact on human health [58,22,31,12]. Information on this diversity and its prevalence can help defining needs in terms of disease surveillance and diagnostic. Moreover, this knowledge can allow to understand virus spread within and between countries. In this context, the interest of our results is double fold. First, we provide support for the interest of shotgun metagenomics in surveys of mosquito-borne viruses in a country with a high viral diversity. Secondly, the detection of SINV opens the way to studies on the prevalence of SINV infection in humans in Burkina Faso and other countries in the West African region.

## Acknowledgments

This work is part of the ArboSud project funded by the 2018 call of the Montpellier University of Excellence programme (MUSE). D. Kabore acknowledges funding from European Union’s Horizon 2020 research and innovation programme (Infravec2 project, grant agreement 731060).

## Supporting information

**Table S1.**
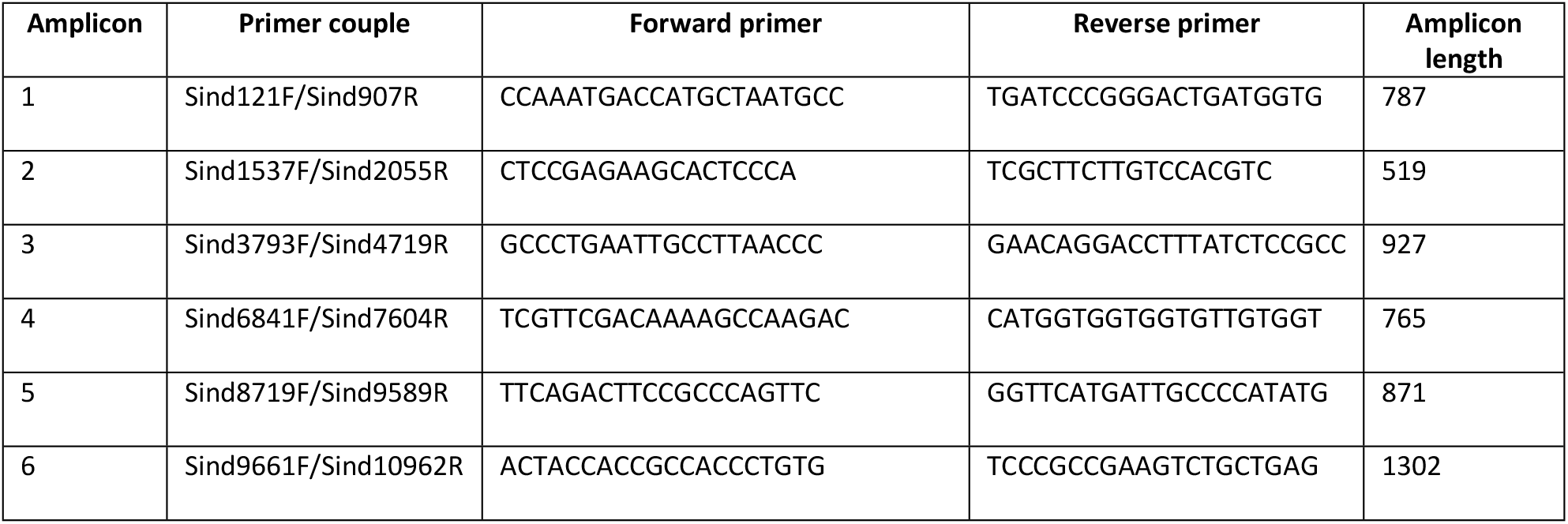
Primers used to amplify and sequence gaps in the SINV genome.

